# Reactive/Non-cooperative individuals advance Population’s Synchronization: Modeling of *Dictyostelium discoideum* Concerted Signaling during Aggregation Phase

**DOI:** 10.1101/2021.05.22.445258

**Authors:** Zahra Eidi, Najme Khorasani, Mehdi Sadeghi

## Abstract

Orchestrated chemical signaling of single cells sounds to be a linchpin of emerging organization and multicellular life form. The social amoeba *Dictiostelium discoiudium* is a well-studied model organism to explore overall pictures of grouped behavior in developmental biology. The chemical waves secreted by aggregating *Dictiostelium* is a superb example of pattern formation. The waves are either circular or spiral in shape, according to the incremental population density of a self-aggregating community of individuals. Here, we revisit the spatiotemporal patterns that appear in an excitable medium due to synchronization of randomly firing individuals, but with a more parsimonies attitude. According to our model, a fraction of these individuals is refusal to amplify external stimulants. Our simulations indicate that the cells enhance the system’s asymmetry and as a result, nucleate early sustainable spiral territory zones, provided that their relative population does not exceed a tolerable threshold.

## Introduction

Cell communication has marked a major transition in the evolution process of life complexity resulted in multicellular organisms contained many cells acting in concert [1, 2]. This ability is coordinated by biochemical signaling networks within individual cells. This process becomes feasible via chemical signaling between fellow creatures. Once the same chemicals are being sensed and produced, every single cell in the population attunes itself to the presence and activities of other cells around it. This collective behavior can lead to the “emergence” of constructions, a new structure only created by a lot of individual interactions between single cells [3]. This is one of the most fascinating and mysterious features of evolutionary life. In particular, collective migration of cohesive cell groups is substantial in organizational constructions like embryogenesis, tissue formation, wound healing ,and cancer invasion [4]. In addition, this collective behavior of unicellulars has been appealing to inspire the engineering of swarm robotics in recent years [5]. All the above complex emergent behaviors arise from a relatively simple behavior of individual entities following a certain ruleset of interactions. The subsequent order and unification appear after that the population density of signaling cells meet a threshold. It turns out that “more is different”, but how many more cells are in need to happen synchronization among the population and global signal propagation? Any answer to this question also might relate to the important ambivalence in the field of developmental biology that which one is precedent over the other: organization or differentiation?

One of the well-established model organisms in developmental biology to address general views of communication, collective behavior, morphogenesis ,and differentiation is the amoeba Dictyostelium discoideum. The amoeba is studied particularly as a borderline between one cell type and two. The cells are capable to evolve from a unicellular to a multicellular organism during their life-cycle by resorting to a chemical mechanism of intercellular communication [6]. During most of its life-cycle, the organism lives in the soil as a single amoeba and feeds on bacteria. When the nutrients are depleted, the cells secrete a chemical called cyclic adenosine monophosphate (cAMP) in response to the pressure. This starts a multicellular developmental process [7]. Cells sense gradients of cAMP which has been emitted from their fellow creatures and direct their chemotactic movements towards regions of its higher concentration. They simultaneously amplify the environmental cAMP concentration by secreting it in their turn. The amplification dynamics results in a periodic cAMP wave propagation process in the medium. The waves propagate from the aggregation center outwardly and guide the chemotactic movement of other cells towards the locus. As a result, up to a million amoeboid cells stream towards the aggregation centers. Thereafter the system goes through a sequence of morphogenetic phases which ultimately land up a fruiting body consisting of two cell types: a delicate stalk of millimeter height, in which about 20% of the cells die to lift the mass of encapsulated spore cells up and hold them off the ground for optimal spore dispersal [8].

The biochemistry behind the process of signaling is acceptably well-understood [9]. The process starts after that the extracellular cAMP bounded to the membrane’s peripheral receptors. This triggers a chain of intracellular chemical reactions which leads to the production afresh of cAMP and its release into the extracellular medium. The cycle attains its terminus after degradation of cAMP by the enzyme phosphodiesterase, through which the receptors get free again [10]. In order to form the signal spatiotemporal patterns, it is necessary that the concentration of re-produced cAMP exceeds that of initially bounded to receptors. This happens through a non-linear autocatalytic process and thus comes at an expensive price [11]. However, during the aggregation process, each and every single cell of the population is confronted with the limits posed by the energetic costs of cellular metabolism for signaling and locomotion while it is in the starving phase of its life cycle [12]. Hence, naturally, they should be inclined to parsimoniously spend their energy, both at the individual and population levels. Although it is possible that the whole number of cells cooperate in signal relaying dynamics throughout the medium, it is a reasonable question to ask that if it is plausible for them to do so as energy-consuming units?

On the other hand, one of the crucial issues in the genesis of global coherent spatiotemporal patterns in a medium of randomly distributed signaling individuals is that how and why the system selects between circular and spiral waves to propagate chemical information from one point to another [13]. Circular waves generally appear as coherent activity of a group of individuals, which are referred as pacemakers. The cells spontaneously oscillate and send out periodic pulses of cAMP [14]. However, spiral waves emerge due to breaking the symmetry of interfacing circular patterns [15]. They are sustainable and not extinguished once they arise. Besides, the oscillation frequency of spiral waves is often higher than circular ones [14, 16]. In principle, the formation of spiral patterns provides a sustainable cue for the individuals to steer them toward a focal point [17]. Analysis of emerging cAMP waves during aggregation process of Dictyostelium shows that, while in areas with low population densities circular patterns are more favored, in zones with higher population densities spiral waves are preferred more [7, 13, 18, 19]; however, there are deviations from this observation [9, 20].

In this study, we use excitable medium formalism as a general framework to elucidate general features of spatiotemporal patterns emerging from cell-cell signaling, plus an extra condition permitting the appearance of population bimodality. The basic idea of the model is an ansatz, which assumes that it is probably more affordable for the population if a fraction of cells does not contribute to amplifying cAMP oscillations in the streaming aggregates and retain their energy for future population survival. We see that this assumption not only is profitable to spend the population’s energy but also advances synchronization of individuals and as a result, earlier emerging sustainable spiral patterns, even in low population densities.

## Materials and Method

In general, an excitable medium is comprised of a continuous set of coupled locally excitable regions, .i.e. cells, which can be both independently stimulated and inhibited. Each cell is characterized by the level of its local extracellular cAMP *u* and the state of its cAMP receptors *υ*. The medium is capable of transmitting information (u value) via promoting spatiotemporal cAMP wave patterns within itself [21]. Components of this medium enjoy of having two distinguishable states: rest and excited. An individual’s rest state provoked to the excited state, provided that the concentration of diffusive stimulus secreted by its neighbors exceeds a threshold. The cell plays the same excitatory role for its adjacent cells in turn and this process initiates a propagating excitation wave throughout the medium. The main dynamical features of a broad class of excitable media can be simulated by a two-component reaction-diffusion system of the form [22]

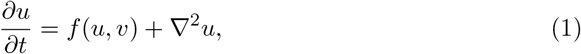

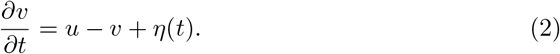

in which the local kinetics of the activator *u* and the inhibitor *υ* is governed by the nonlinear functions *f*(*u, υ*) in Eq. 1 and the linear function *u* − *υ* in Eq. 2, respectively. We assume that *f*(*u, υ*) = *ϵ*^−1^*u*(1 − *u*)(*u* − *u*_*th*_), where *u*_*th*_ = (*υ* + *b*)/*a*. The parameter *ϵ* indicates excitability of the system which is the time-scale difference between the fast excitatory variable *u* and the slow refractory variable *υ* while *a* and *b* are the controlling parameters of the model. Our numerical simulations are based on a slightly modified version of Barkley model [23]. In our model, we assume that a given cell is in its excited state if its corresponding value of *u* exceeds the size of boundary layer *δ*, See Fig. 1-A. In Eq. 2, the function *η*(*t*) reflects affected noise on the inhibitor variable *υ*. We assume that *η* is driven by a colored noise known as Ornstein–Uhlenbeck process with 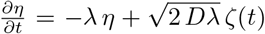, which has intensity *D* and correlation time *λ*^−1^. Here, *ζ*(*t*) is Gaussian white noise of mean zero and correlation function < *ζ*(*t*)*ζ*(*s*) >= *δ*(*t* − *s*) [24].

The local dynamics of an excitable medium, that is the dynamics in the absence of diffusion, See Fig. 1-A, is characterized by a stable but excitable fixed point. The point sits at the intersection locus of *u* and *υ* nullclines, which are the answers of *f* (*u, υ*) = 0, and *u − υ* = 0 line respectively. Fig. 1-A provides a schematic illustration of typical corresponding nullclines. Small perturbations of the rest state decay and return to the fixed point immediately. However, perturbations that exceed the boundary layer size *δ* increase and decay only after the system has performed a large excursion in the *uυ*-phase plane. In this case, after passing a long loop the system relaxes to the fixed point and stands by for the next itinerary.

**Fig 1.**
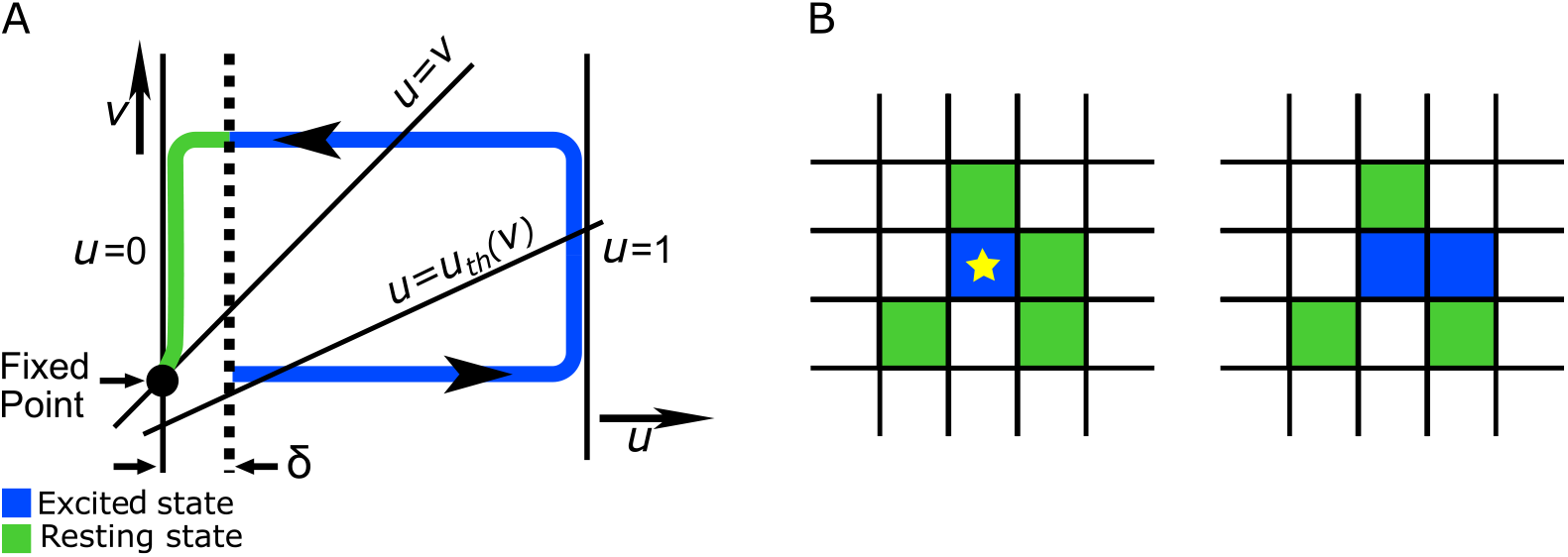
(Color online) Schematic illustration of local dynamics. Corresponding dynamics of the excitable system in *uυ*-plane is depicted by its nullclines. The *u* nullcline includes three separate lines: *u* = 0, *u* = 1 and *u* = (*υ* + *b*)*/a*, while the line *u* = *υ* is the *υ* nullcline. The dotted line indicates the boundary layer *δ* which all the initial conditions within it decay to the fixed point. The itinerary in the *uυ*- plane starts haphazardly once the value of *u* exceeds the amount of *δ*. Eventually, the itinerary returns to the fixed point after passing a large excursion, adapted from [23] (A). The medium consists of excitable cells spatially connected to each other by a diffusion-like coupling. An exited cell (the blue asterisk unite) is capable of triggering excitation through its neighbors with the help of diffusing cAMP molecules (B).

A special feature of a spatially extended system of excitable media is that such systems promote local impulses i.e. leading to propagating spatiotemporal patterns throughout the system. In reality, this takes place provided that the diffusive coupling of the excitable elements is of adequate strength. The diffusive mechanism with which adjacent excitatory cells stimulate each other via signaling messenger *u* is schematically illustrated in Fig. 1-B. Here the blue asterisked unit is in its excited state while all its nearby cells relax in their rest state,.i.e. they are green. The strength of the diffusion of the local *u* is sufficient to impel one of at rest neighbors cell, say its right neighbor, into its excited state, as depicted in Fig. 1-B. The excited state of the blue cell soon becomes refractory, a state in which the cell neither stimulates nor is stimulated by its neighbors. Note that the cells have to be close enough so that diffusion of signaling messenger works constructively to share the information of the amount of *u* from one cell to the other.

As a matter of fact, the nonlinear cubic term *f* (*u, υ*) in Eq. 1 is necessary to shape patterns. An evident feature in the aggregation of Dictyostelium amoebae is that the environmental cAMP signals are amplified by the individuals [6]. But, what if there are a fraction of the individuals which are refusal to amplify external stimulants? Let us suppose that the population of aggregating cells is not pure, but mixed including two types based on their tendency to treat proactive/cooperative or reactive/non-cooperative behavior in the population. A group of cells are capable (and tend) to boost the external cAMP signaling messenger via secreting more cAMP locally; This way, they form pacemakers that trigger periodic pulses of cAMP in the medium. This is modeled by nonlinear term in the Eq. 1. We refer to this group of individuals as proactive/cooperative cells. The other type of cells is potentially unable to amplify the environmental cAMP concentration. Thus, they are reactive/non-cooperative. These individuals align their locomotion toward the gradient of the chemical stimulus, utilizing the signal released by other cells. Based on the model, the same set of equations as Eq. 1 governs the dynamics of reactive/non-cooperative cells except that instead of the cubic term in this equation, they possess a linear term as *u* − *u*_*th*_. Now, the question is that, considering the energy cost of nonlinear behavior and time limitations during which intercellular biochemical reactions occur to form spatiotemporal patterns, what is the ideal combination of the cells in the mixed population? In the other words, assuming that the population density of the whole population is *ϕ*_1_ and among these cells, the fraction of *ϕ*_2_ are reactive/non-cooperative ones, what is the optimum value of *ϕ*_2_ by which the system can bear the subsistence of the cells before depriving spatiotemporal patterns totally?

## Results

### Population density of individuals has to meet a threshold for global spatiotemporal patterns appearance

Circular and spiral waves are two generic spatiotemporal patterns of excitable systems. Depending on the system’s initial condition and interacting parameters, the spatial diversity distribution of individuals and frequencies of oscillatory elements the corresponding features of circular waves or spiral ones are favored by the medium [19]. The aim of current investigation is to study the probable presence of a mixed population at the early stages of cell aggregation. Accordingly, we restrict the simulations only to the wave propagation properties upon which even in high densities of a single species population the shape of patterns are circular. For a thorough list of parameters see section. Shown in Fig. 2 are spatiotemporal snapshots of different lattices exclusively composed of proactive/cooperative cells with different population densities *ϕ*_1_. From Fig. 2 , this is apparent that for low pure population densities (*ϕ*_1_ < 0.4), the local chemoattractant cAMP concentration is insufficient to synchronize the individuals in a way that global territory zones of patterns form. Throughout this study, we focus our attention on the system whose at least *ϕ*_1_ = 0.4 fraction of its lattice units are covered with cells.

**Fig 2.**
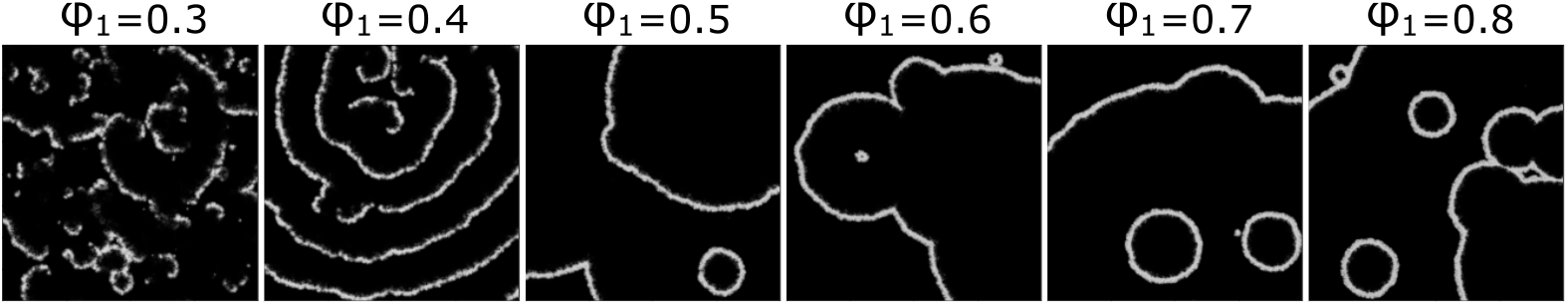
Emergent patterns on different lattices including randomly distributed firing cells with variant *ϕ*_1_. *ϕ*_1_ is population density of the cells which determines the number density of occupied units on the lattices of size 400 × 400. Here, the cells are exclusively of proactive/cooperative type. All parameters of the model are kept the same for these systems and listed in section.

### Spiral waves appear in a field composed of the mixed population even at low population densities

One of the most striking features of non-pure bimodal populations is the emergence of spiral waves at low population densities. Fig. 3 panels: A, D, G, J, M shows time-lapse frames of spatiotemporal patterns appearing in systems with different *ϕ*_2_ values, where just half of the lattice units are occupied,. i.e. population density of all systems are *ϕ*_1_ = 0.5. Fig. 3. A illustrates appearing and disappearing circular waves where the whole cells are proactive ones (*ϕ*_2_ = 0) and panels D, G, J ,and M show signaling patterns of systems in which population density of reactive/non-cooperative individuals are none-zero with the *ϕ*_2_ value indicated beside each row, See also Supplemental Video 1 to Video 5. The difference of these patterns becomes exquisite when a small fraction of cells (say *ϕ*_2_ = 0.1) are of reactive/non-cooperative type, Fig. 3-D, where spiral patterns emerge, Supplemental Video 2. Indeed, it seems in bimodal populations with a low population density of reactive cells, circular waves interact with one another setting up into spiral waves. However, when the reactive population density exceeds the threshold value of *ϕ*_2_ = 0.1, the waves extinguish just to fluctuate in local cAMP concentration and the distinguishable spiral waves disappear gradually.

**Fig 3.**
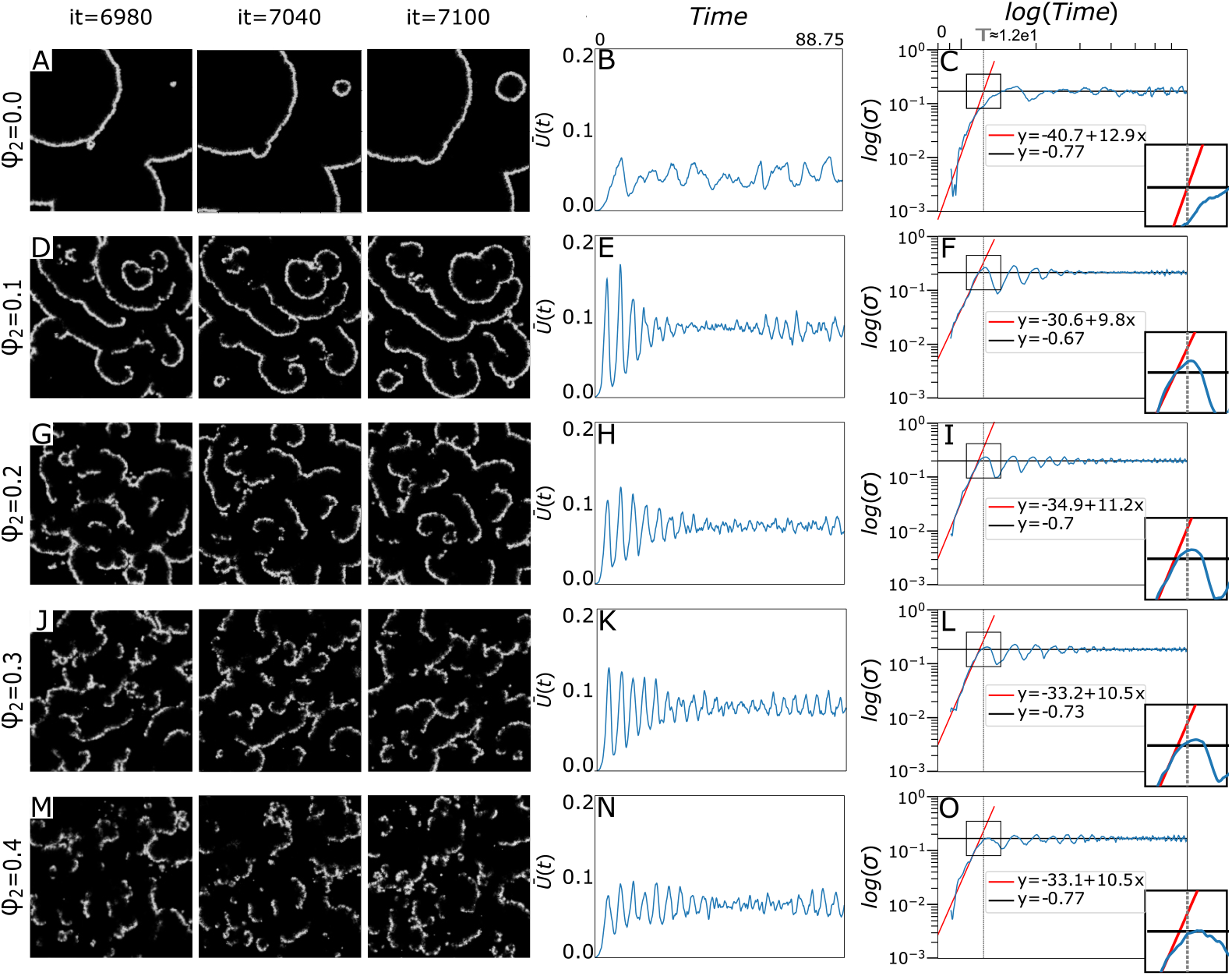
(Color online) Propagating wave through a system with *ϕ*_1_ = 0.5 and *ϕ*_2_ = 0.0 (A), *ϕ*_1_ = 0.5 and *ϕ*_2_ = 0.1 (D), *ϕ*_1_ = 0.5 and *ϕ*_2_ = 0.2 (G), *ϕ*_1_ = 0.5 and *ϕ*_2_ = 0.3 (J) and *ϕ*_1_ = and *ϕ*_2_ = 0.4 (M). There are 60 iteration interval between the illustrated subsequent patterns, here ‘it’ is an abbreviation for iteration. Emerging spiral waves are distinguishable only at (D) See also Supplemental Video 1 to Video 5. Mean concentration of signaling agent, calculated as 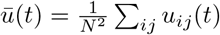 where *u*_*ij*_ (*t*) is the cAMP concentration of point (*i, j*) at time *t*, in a system with *ϕ*_1_ = 0.5 and *ϕ*_2_ = 0.0 (B), *ϕ*_1_ = 0.5 and *ϕ*_2_ = 0.1 (E), *ϕ*_1_ = 0.5 and *ϕ*_2_ = 0.2 (H), *ϕ*_1_ = 0.5 and *ϕ*_2_ = 0.3 (K) and *ϕ*_1_ = 0.5 and *ϕ*_2_ = 0.4 (N). Logarithmic scale of standard deviation of *u*, calculated as 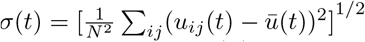, for a system with *ϕ*_1_ = 0.5 and *ϕ*_2_ = 0.0 (C), *ϕ*_1_ = 0.5 and *ϕ*_2_ = 0.1 (F), *ϕ*_1_ = 0.5 and *ϕ*_2_ = 0.2 (I), *ϕ*_1_ = 0.5 and *ϕ*_2_ = 0.3 (L) and *ϕ*_1_ = 0.5 and *ϕ*_2_ = 0.4 (O). Red and black lines are representatives of growing phase and oscilatory one, respectively. In each plot red line is the linear regression of log(*σ*(*t*)) for the first 300 iterations of every simulation. Black line in each diagram is the mean value of log(*σ*(*t*)) in the last 3000 iterations in each simulation. The magnified window located in right bottom of panels C, F, I, L and O illustrates the crossover time of the system dynamics between these two regimes. The vertical gray dashed line corresponds to the crossover time in a pure system with *ϕ*_1_ = 0.5 and *ϕ*_2_ = 0.0 which is the intersection point of black and red lines in panel (C).

To assess the concentration level of signaling messenger *u*, we compute the mean-field concentration of the stimulant in the medium as 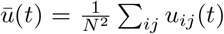 where *u*_*ij*_ (*t*) is the cAMP concentration of point (*i, j*) at time *t*. It seems that even in the presence of the reactive/non-cooperative cells the average concentration of stimulants remains intact, between 0.05 and 0.08, Compare trend value of 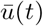 plot in panels: B, E, H, K, and N of Fig. 3. This brings us to the conclusion that the situation of the system has not change for the worse in this case after making the population bimodal. The concentration standard deviation, which is the root mean squared fluctuation in the cAMP chemical concentration reads as 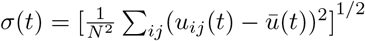. Comparison of panels C, L, I, F, and O of Fig. 3 persuade us that the behavior of time evolution of stimulant concentration standard deviation is almost identical in each case. It has typically two regions separated by a crossover time, which is the intersection point of two lines defining two distinct dynamical behavior. The concentration width initiates by growing exponentially and after passing the crossover time, its dynamics follows an oscillating behavior around some saturated value. The saturated regime occurs during the very time period that the spatiotemporal waves are observable in the medium. In Fig. 3, the crossover time of the system which only includes proactive/cooperative cells, *T*, is illustrated as a dashed line. The line is considered as a benchmark against which one can decide about the same quantity in other systems and inspect the impact of the presence of reactive/non-cooperative cells on the quantity. Remarkably, the crossover time between two growing and oscillating regimes occurs earlier when reactive/non-cooperative cells attend in the medium. One may conclude that reactive/non-cooperative individuals advance the population’s synchronization to construct spatiotemporal patterns.

The growth slope of the plots before crossover time is indicated inside each panel, see panels C, L, I, F, and O of Fig. 3. The quantity is a measure of how fast the fluctuating dynamics of concentration evolve to reach out the oscillatory dynamics. It seems that the slope has descending behavior with respect to the presence of reactive individuals.

### Mere presence of reactive/non-cooperative individuals do not affect population’s synchronicity markedly

As an important illustration of the general features of collective signalling,, we consider the cross-correlation *S* for the *u* variable as space and time-averaged nearest-neighbor distance of all elements, normalized by the total spatial amplitude variance 25]. T equantity measures the coherency of the patterns and is defined as 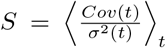 where *Cov*(*t*) spat l auto-covariance of nearest neighbors at time *t* elucidated as 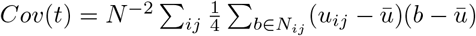. Here, *b* takes up *u* values of all 4 elements of a von Neumann neighborhood *N*_*ij*_ at each lattice site and the bracket ⟨⟩_*t*_ denotes averaging over the total integration time *t*. This is a quantitative measure for the relative change of the order in a spatially extended system. Its maximal value is *S* = 1 for a completely synchronized network in space and time. Table 1 provides details about the *S* value for variant pairs of reactive and proactive mixture in a medium in which half of its surface is covered with the cells. The three rows of the table represent the *S* value of three independent realizations for each simulated system. Comparing the first array of each row, which belongs to a system in the absence of reactive individuals, with other ones in each array, we see that the *S* value experiences a modest decline (around 0.01) in presence of reactive cells. However, this quantity hardly changes and remains fairly constant when the fraction of reactive cells increases in the medium. The survey of the value for a bunch of systems with variant pairs of *ϕ*_1_ and *ϕ*_2_ reveals that the *S* value for the whole systems are between 0.930 and 0.950.

**Table 1.** Comparison of *S* values of the media with different proportions of reactive/non-cooperative cells whose surface is covered with randomly distributed firing cells with population density *ϕ*_1_ = 1.

| | $\phi_2 = 0$ | $\phi_2 = 0.1$ | $\phi_2 = 0.2$ | $\phi_2 = 0.3$ | $\phi_2 = 0.4$ |
| --- | --- | --- | --- | --- | --- |
| **S** | 0.949 | 0.940 | 0.940 | 0.940 | 0.938 |
|  | 0.949 | 0.939 | 0.939 | 0.939 | 0.938 |
|  | 0.950 | 0.940 | 0.940 | 0.938 | 0.938 |

### Low proportions of reactive/non-cooperative individuals in a population contributes to the formation of sustainable spiral waves

Fig. 4 illustrates an arranged snapshots of a dozen mixed populations with different pairs of *ϕ*_1_ and *ϕ*_2_, which are addressed on vertical and horizontal margins, respectively. The disposition of snaps in the array suggests that not only arising circular or spiral patterns are not erratic, but also it can provide a methodical ordering to predict the related pattern of every pair of population mixture. As depicted in Fig. 4 in pure populations (with *ϕ*_1_ ≥ 0.4) with parameters listed in section circular waves are favored. This category is marked with a blue boundary in the figure. It is seen that the signals in the pure system with *ϕ*_1_ = 0.3 are diluter than distinguished pattern can emerge. Although the pattern of the pure system with *ϕ*_1_ = 0.4 sounds to be circular, due to apparent differences with other pure systems we excluded it from the blue border. From left to right we have an ascending amount of reactive/non-cooperative cells in the populations and simultaneously, a descending order in the patterns. In the territory of the red border in which a low fraction of cells is of reactive/non-cooperative type spiral waves appear. It seems that low amounts of these cells in the medium enhances the system asymmetry and promote the early nucleation of spiral forms. However, the system can tolerate subsistence of these cells only up to a threshold depending on the population density of whole cells, .e.g, the threshold is around *ϕ*_2_ = 0.2 in the system with *ϕ*_1_ = 0.8. When the relative number of these cells increases in the population the patterns become more inconspicuous gradually until only a fine mist of dilute signals come into sight at the right side of the array, see the area restricted with yellow and orange boundary lines in Fig. 4).

**Fig 4.**
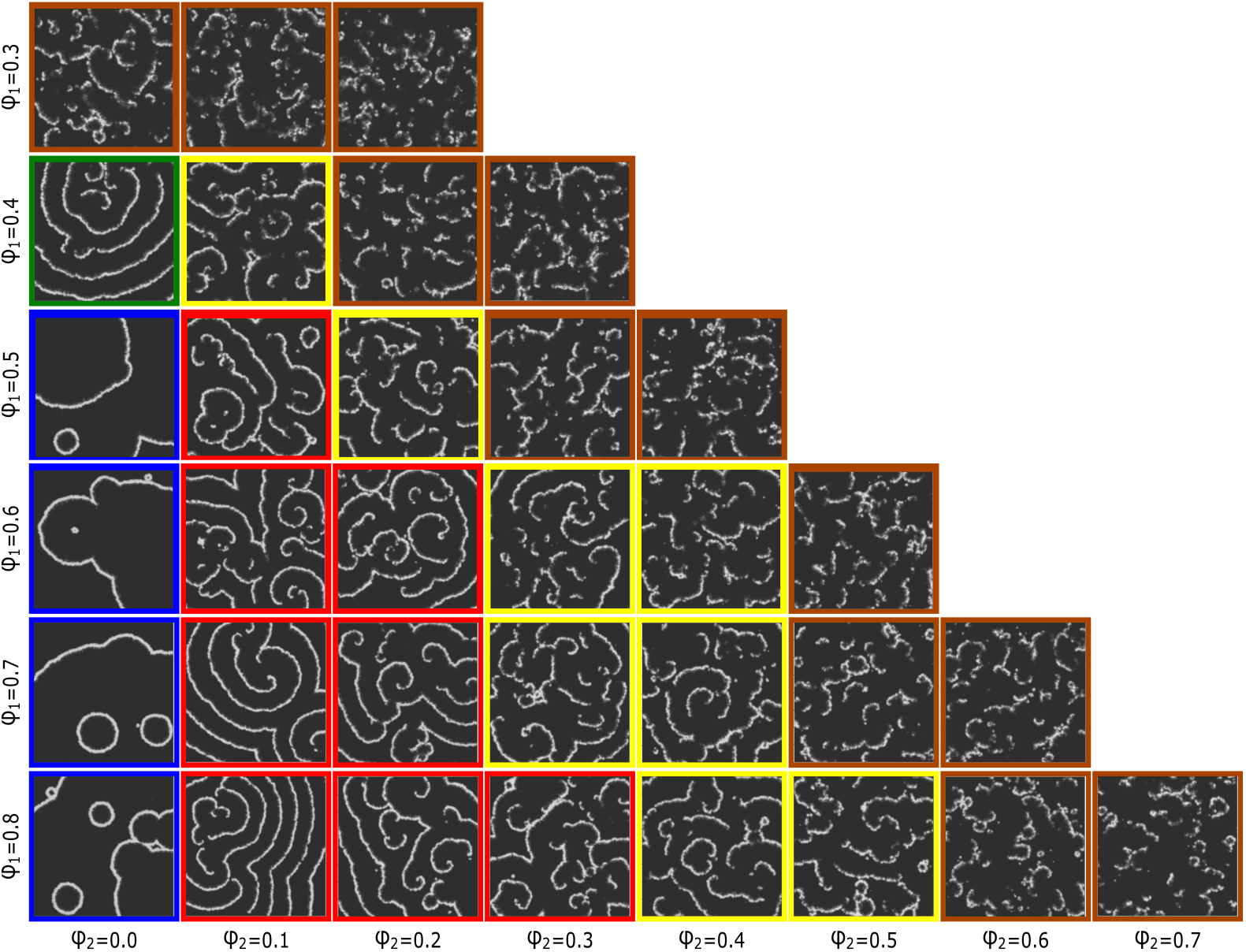
(Color online) Disposition of pattern snapshots on a common instance of time in lattices of 400 × 400 size, covered with mixed populations with different pairs of *ϕ*_1_ and *ϕ*_2_. *ϕ*_1_ is the population density of the whole cells on each lattice and *ϕ*_2_ is the fraction of reactive/non-cooperative cells among the population. The related row and column of each pattern in the array are appointed by their *ϕ*_1_ and *ϕ*_2_ indicated in vertical and horizontal margins, respectively. For example, the bottom row includes the systems with the same population density of individuals *ϕ*_1_ = 0.8 but with a variant fraction of reactive/non-cooperative ones *ϕ*_2_. Similar patterns are enclosed with a colored border. Pure systems are arranged in the first column.

## Discussion

The oscillatory signals during the self-aggregation period of *Dictyostelium discoideum* are reinforced through two major processes: intracellular biochemical feedback loops that make the cells relay the external signals, and cell recruitments in response to the signals which, in turn, leads to configurations with more efficacy in signaling [3]. Here, we merely concentrate on the first process and adjourn the second one to the next studies. In the current study, we implemented a parsimonious attitude to model a concerted signaling process through a mixed population of randomly distributed excitable coupled individuals. Based on the model, the population is composed of two types of cells: Proactive/cooperative cells which contribute toward relaying external signaling messenger of cAMP, and reactive/non-cooperative ones that refuse to relay the environmental cAMP. Population bimodality can emerge even when biological processes are homogenous at the cell level and the environment is kept constant [26]. There are pieces of evidence that cheating occurs in mixed populations of wild clones [3, 27]. Although selfish behavior known as cheating is reported in the slug stage, one might track back its trace to prior stages of social communication and interaction between individual units that are spatially separated, as well [6].

Experiments show that in non-living reaction-diffusion systems such as the Belouzov–Zhabot reaction, oscillators often form around inhomogeneities [28]. In contrast, evolving systems covered by populations of Dictyostelium are intrinsically heterogeneous. Their variant kinetic properties during the course of time have a profound impact on waveform competition [15]. Noise and variability in the form of cell-to-cell differences are common themes in the study of biological organizations. Essentially, instead of suppressing or filtering out the noise and eliminating diversity, individual living matters often exploit these features to drive many of theirs intracellular and extracellular processes [3, 29]. Differences between pattern formation in biological and chemical or physical systems are attributed to these properties [29]. Principally, self-organized physical systems are consist of functionally identical elemental parts, while components in that of biological systems are diverse in their essence [30]. Recently, it has shown that stable activity centers appear spontaneously in areas of higher cell density with the oscillation frequency of these centers depending on their density [18]. On the other hand, it has proved that diversity-induced resonance has a principal role in pattern competition between circular and spiral waves [31]. In particular, the increase and decrease of the spiral wave count with increasing diversity are statistically regulated by the number of oscillatory elements in the lattice, rather than by the frequency differences between target and spiral waves [19]. In our model, the individuals essentially enjoy of an inherent inhomogeneity due to the bimodal nature of the population, i.e. composed of reactive/non-cooperative cells and proactive/cooperative ones. As it is evident in Fig. 2 circular wave frequencies in different systems covered only by proactive/cooperative individuals are identical. Accordingly, after substituting a fraction of these cells with reactive ones, one can attribute the emerging patterns of spirals to the intrinsic inhomogeneity of cells, especially at low population densities, see Fig. 3. This consequence is likely to be of help to explain why for a particular strain and circumstance one pattern tends to dominate. This is an important question because they bear on early events governing the self-aggregation of the slime mold organizational process. Any answer to this question might shed indirect light on the classic question that: Which one comes first? Differentiation or Morphogenesis. From experimental insight, it is traditionally viable to measure the amount of local concentration of cAMP relay by light scattering waves caused by the synchronized cell movements at variant developmental stages of Dictyostelium aggregation [32–34]. However, there are reports that show no clear evidence exists for cAMP relay organizing collective cell migration at multicellular stages [35]. Most recent experiments report cAMP waves in living cell populations using cAMP specific fluorescence resonance energy transfer (FRET) techniques during aggregation process [7, 36]. On the other hand, Hashimura, et.al challenged the traditional view about the role of cAMP relay in the organization of collective cell migration recently [37].

In our in silico experiments, apparent resemblances of emerging patterns in different systems are deemed to classify them, see Fig. 4. To abandon ourselves from the likely occurring errors of manual monitoring, it is necessary to develop an automated machine learning-based method of data analysis and image processing to compare the patterns and set them out in Fig. 4. Nevertheless, it sounds that 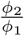 reckoned to be a relevant quantity in apparent similarities between the snapshots of different systems. Another relevant quantity in shared features of patterns in the presence and absence of reactive/non-cooperative cells is the growth incline of cAMP concentration standard deviation, panel *C* in comparison with panels *F*, *I*, *L*, and *O*. The apparent diminution of the slope of red lines in the presence of reactive individuals reminds us of different methods of deposition of colloidal aggregates resulting in vastly different exponential growth slopes of the width of mean height, where the slopes define universality classes of roughening processes [38]. Certain statement about this requires a thorough survey and would be a potentially interesting area for future research.

## APPENDIX: Numerical simulation and data analysis

As mentioned above, we perform numerical experiments using the slightly modified Barkley model [23]. We simulate the system of Eq. 1 using the finite difference method:

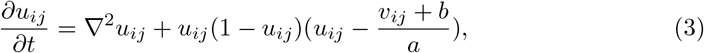

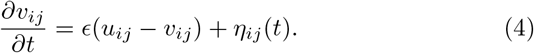

As illustrated in Fig. 1-A, the boundary layer with size *δ* defines the threshold value for *u*_*ij*_.We assume that *u* = 0 within the interval [0.*δ*),. i. e. in every cell whenever *u*_*ij*_ < *δ*, its value in the next update will be considered equals to zero. The values of the variables are updated using a discrete-time derivative Euler method. Here, we compute ∇^2^ as the discrete Laplace operator of a 2*D* variable on the grid, using a five-point stencil finite difference method as 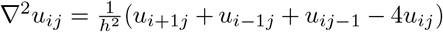 where *h* is the grid spacing. We perform our simulations on a 400 × 400 lattice with Neumann boundary condition state that the spatial derivatives with respect to the normal vectors are null on the boundaries of the domain. We implement these boundary conditions by duplicating values in matrices *u* and *υ* on the edges at each time step. Throughout this study, the physical values are set as *a* = 0.3, *b* = 0.01 *and ϵ* = 0.005. Besides, numerical values are adjusted as *h* = 0.25, Δ*t* = 0.05 *and δ* = 0.0125. The parameters are regulated to yield the network in a sub-excitable state, i.e., the formation of sustainable spatial patterns is impossible without the existence of *η* term. Parallel with the integration of the deterministic part, we implement the Euler–Maruyama technique with a time step Δ*t* to integrate the *η*(*t*) stochastic dynamics,

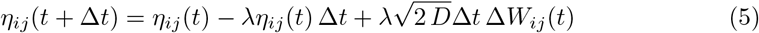

where the Δ*W* (*t*) are independent Gaussian random numbers with unit variance (known as increments of the Wiener process [24]). Stochastic values are set as *D* = 1 and *λ* = 1*/*300. Considering them as the transient period, signals occurring in the first 900 time steps are excluded from the calculations. Besides, the edges of the lattice is removed in the analysis to avoid artifacts from the event recognition at the borders.

## Notes

### Competing Interest Statement

The authors have declared no competing interest.

## References

1. Herron MD, Michod RE. Evolution of complexity in the volvocine algae: Transitions in individuality through Darwin’s eye. Evolution. 2008; 62 :436–451.

2. Queller DC. Relatedness and the fraternal major transitions. Philos Trans R Soc Lond B Biol Sci. 2000;355:1647–1655.

3. Reid CR, Latty T. Collective behaviour and swarm intelligence in slime moulds. FEMS Microbiology Reviews. 2016 Nov;40(6):798–806.

4. Friedl P, Gilmour D. Collective cell migration in morphogenesis, regeneration and cancer. Nat. Rev. Mol. Cell Biol. 2009;10:445–457.

5. Parhizkar M, Serugendo GM. Agent-based models for first- and second-order emergent collective behaviors of social amoeba Dictyostelium discoideum aggregation and migration phases. Artificial Life and Robotics Journal. 2018;23(4): 498–507.

6. Kessin RH. Dictyostelium: evolution, cell biology, and the development of multicellularity. Cambridge University Press; 2001.

7. Singer G, Araki T, and Weijer CJ. Oscillatory cAMP cell-cell signalling persists during multicellular Dictyostelium development. Commun. Biol. 2019;2:139.

8. Brock DA, Gomer RH. A cell-counting factor regulating structure size in Dictyostelium. Genes Dev.1999;13(15):1960–1969.

9. Pálsson E, Cox EC. Origin and evolution of circular waves and spirals in Dictyostelium discoideum territories. Proc. Natl Acad. Sci. USA.1996 Feb 6;93(3):1151–5. doi: 10.1073/pnas.93.3.1151.

10. Iglesias PA. Feedback control in intracellular signaling pathways: Regulating chemotaxis in Dictyostelium discoideum. European Journal of Control. 2003;9(2-3):227–36.

11. S. K. Scott, Chemical Chaos. Oxford University Press, Oxford; 1991.

12. Strassmann JE, Queller DC. Evolution of cooperation and control of cheating in a social microbe. Proc. Natll. Acad. Sci.2011;108(2):10855–10862.

13. Lee KJ, Goldstein RE, Cox EC. cAMP waves in Dictyostelium territories. Nonlinearity. 2002; 15: C1–C5.

14. Gregor T, Fujimoto K, Masaki N, Sawai S. The onset of collective behavior in social amoebae. Science. 2010 May;328(5981):1021–5. doi: 10.1126/science.1183415. Epub 2010 Apr 22.

15. Lauzeral J, Halloy J, Goldbeter A. Desynchronization of cells on the developmental path triggers the formation of spiral waves of cAMP during Dictyostelium aggregation Proc. Natl Acad. Sci.1997; 94 (17):9153–9158.

16. Sgro AE, Schwab DJ, Noorbakhsh J, Mestler T, Mehta P, Gregor T. From intracellular signaling to population oscillations: bridging size- and time-scales in collective behavior. Mol Syst Biol. 2015;11:779.

17. Noorbakhsh J, Schwab DJ, Sgro AE, Gregor T, Mehta P. Modeling oscillations and spiral waves in Dictyostelium populations. PHYSICAL REVIEW E. 2015 Jun;91(6):062711.

18. Vidal-Henriquez E, Gholami A. Spontaneous center formation in Dictyostelium discoideum. Scientific Reports. 2019; 9:3935.

19. Grace M, Hütt MT. Pattern competition as a driver of diversity-induced resonance.Eur. Phys. J. B. 2014;87,29.

20. Durston AJ. Dictyostelium discoideum aggregation fields as excitable media. J. Theor. Biol. 1973;42: 483–504.

21. Sager BM. Propagation of traveling waves in excitable media. Genes Dev. 1996;10:2237–2250.

22. Murray JD. Mathematical Biology II, Springer; 2003.

23. Barkley D. A model for fast computer simulation of waves in excitable media. Physica D. 1991;49:61–70.

24. Laing C, Lord CJ. Stochastic Methods in Neuroscience, Oxford University Press, New York; 2010.

25. Busch H, Kaiser F. Influence of spatiotemporally correlated noise on structure formation in excitable media. Phys. Rev. E. 2003;67, 041105.

26. Fernandez-de-Cossio-Diaz J, Mulet R, Vazquez A. Cell population heterogeneity driven by stochastic partition and growth optimality. Scientific Reports. 2019; 9,9406.

27. Strassmann JE, Zhu Y, Queller DC. Altruism and social cheating in the social amoeba Dictyostelium discoideum. Nature.2000;408,965–7.

28. Agladze K, Keener JP, M’uller Sc, Panfilov A. Rotating spiral waves created by geometry. Science. 1994;264,1746.

29. Grace M, Hütt MT. Regulation of Spatiotemporal Patterns by Biological Variability: General Principles and Applications to Dictyostelium discoideum. PLOS Computational Biology. 2015;11(11):e1004367.

30. Grace M, Hütt MT. Predictability of spatio-temporal patterns in a lattice of coupled FitzHugh–Nagumo oscillators. J R Soc Interface. 2013 Jan 24;10(81):20121016.

31. Glatt E, Gassel M, Kaiser F. Variability-sustained pattern formation in subexcitable media. PHYSICAL REVIEW E. 2007;75: 026206.

32. Alcantara F, Monk M. Signal propagation during aggregation in the slime mould Dictyostelium discoideum. J. Gen. Microbiol. 1974;85:321–334.

33. Rietdorf J, Siegert F, Weijer CJ. Analysis of optical density wave propagation and cell movement during mound formation in Dictyostelium discoideum. Dev. Biol. 1996;177:427–438.

34. Dormann D, Weijer CJ. Propagating chemoattractant waves coordinate periodic cell movement in Dictyostelium slugs. Development. 2001;128:4535–4543.

35. Wang B, Kuspa A. Dictyostelium development in the absence of cAMP. Science. 1997;277:251–254.

36. Kamino K, Kondo K, Nakajima A, Monda-Kitahara M, Kaneko K, Sawai S. Fold-change detection and scale invariance of cell-cell signaling in social amoeba. Proc. Natl. Acad. Sci. USA. 2017;114:E4149–E4157.

37. Hashimura H, Morimoto YV, Yasui M, Ueda M. Collective cell migration of Dictyostelium without cAMP oscillations at multicellular stages. Commun. Biol. 2019;2,34.

38. Barabasi AL, Stanley HE. Fractal Concepts in Surface Growth. Cambridge University Press; 1995.

